# An inventory of early branch points in microbial phosphonate biosynthesis

**DOI:** 10.1101/2021.04.07.438883

**Authors:** Siwei Li, Geoff P. Horsman

## Abstract

Microbial phosphonate biosynthetic machinery has been identified in ~5% of bacterial genomes and encodes natural products like fosfomycin as well as cell surface decorations. Almost all biological phosphonates originate from the rearrangement of phosphoenolpyruvate (PEP) to phosphonopyruvate (PnPy) catalyzed by PEP mutase (Ppm), and PnPy is often converted to phosphonoacetaldehyde (PnAA) by PnPy decarboxylase (Ppd). Seven enzymes are known or likely to act on either PnPy or PnAA as early branch points en route to diverse biosynthetic outcomes, and these enzymes may be broadly classified into three reaction types: hydride transfer, aminotransfer, and carbon-carbon bond formation. However, the relative abundance of these branch points in microbial phosphonate biosynthesis is unknown. Also unknown is the proportion of *ppm*-containing gene neighborhoods encoding new branch point enzymes and potentially novel phosphonates. In this study we computationally sorted 434 *ppm*-containing gene neighborhoods based on these seven branch point enzymes. Unsurprisingly, the majority (56%) of these pathways encode for production of the common naturally occurring compound 2-aminoethylphosphonate (AEP) or a hydroxylated derivative. The next most abundant genetically encoded intermediates were phosphonoalanine (PnAla, 9.2%), 2-hydroxyethylphosphonate (HEP, 8.5%), and phosphonoacetate (PnAc, 6%). Significantly, about 13% of the gene neighborhoods could not be assigned to any of the seven branch points and may encode novel phosphonates. Sequence similarity network analysis revealed families of unusual gene neighborhoods including possible production of phosphonoacrylate and phosphonofructose, the apparent biosynthetic use of the C-P lyase operon, and a virus-encoded phosphonate. Overall, these results highlight the utility of branch point inventories to identify novel gene neighborhoods and guide future phosphonate discovery efforts.

**IMPACT STATEMENT:** Microbially-produced phosphonates are relatively rare and underexplored but include medically and agriculturally important molecules like fosfomycin and phosphinothricin, respectively. Because a single enzyme called phosphoenolpyruvate mutase (Ppm) inititates almost all phosphonate production, the composition of the ‘gene neighborhood’ surrounding a Ppm-encoding gene can inform hypotheses regarding the chemical output of this chromosomal region. After the initial Ppm-catalyzed reaction there are only a limited set of subsequently acting enzymes, or ‘branch points’, to direct these early-stage phosphonates to alternate chemical fates. However, the relative abundance of different branch points – or the existence of new ones – has not been evaluated. This study provides just such a ‘branch point inventory’ to determine relative proportions of known branch points and assess the diversity within each branch point. Significantly, this study suggests that a significant proportion (~13%) of gene neighborhoods do not fit into known branch points and therefore may be fertile hunting grounds for new phosphonate biochemistry.

**Data Summary:** Supporting information is available at Scholars Portal Dataverse (https://dataverse.scholarsportal.info/) with DOI 10.5683/SP2/T33ZP6. This includes scripts and the network data for visualizing in BiG-SCAPE and Cytoscape.

## INTRODUCTION

Phosphonic and phosphinic acid metabolites contain a carbon-phosphorus bond and represent a commercially successful but underexplored class of biological molecules exemplified by the antibiotic fosfomycin and the herbicide phosphinothricin.[1] In addition to diffusible small molecules, about half of all microbial phosphonate biosynthetic pathways are thought to produce phosphonoglycan and phosphonolipid cell wall constituents, yet almost nothing is known about their biological significance.[2] Moreover, phosphonate metabolism continues to unveil new and surprising catalytic transformations with strong potential to impact biocatalytic production of fine chemicals.[3, 4]

Almost all known biological phosphonates originate from rearrangement of phosphoenolpyruvate (PEP) to phosphonopyruvate (PnPy) catalyzed by phosphoenolpyruvate mutase (Ppm) (**Figure 1**).[5] Detecting Ppm activity long eluded researchers because PEP is heavily favored at equilibrium, and most phosphonate biosynthetic pathways couple the activity of Ppm with that of a second irreversible decarboxylation to phosphonoacetaldehyde (PnAA) catalyzed by PnPy decarboxylase (Ppd).[6] A typical third step involves formation of the most common phosphonate metabolite 2-aminoethylphosphonate (AEP) by the action of AEP transaminase (AEPT).[7]

**Figure 1.**
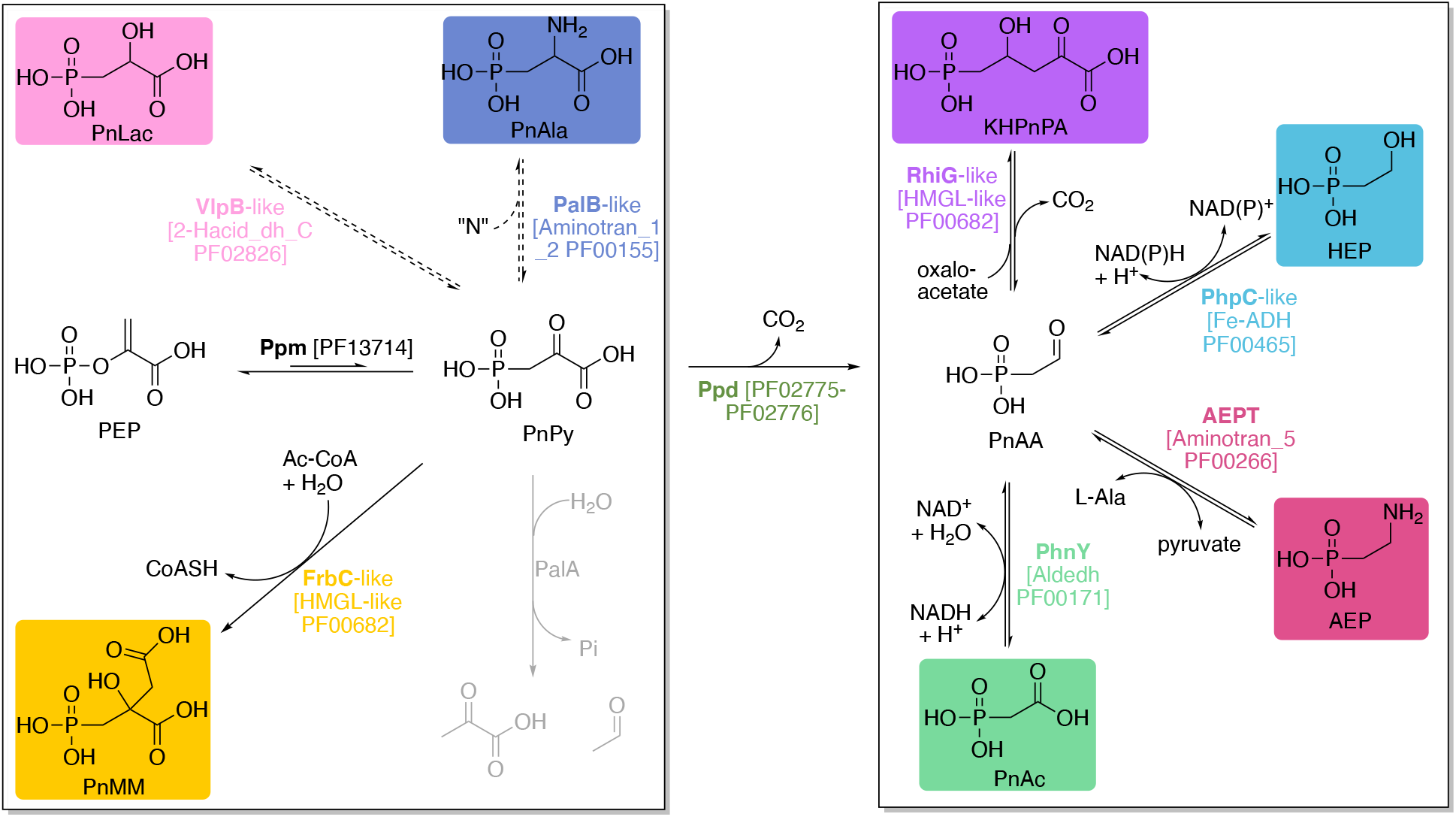
Overview of early branch points in phosphonate biosynthesis. **Left side:** Pre-Ppd transformations. Phosphonate biosynthesis starts from Ppm-catalyzed rearrangement of PEP to PnPy, which is followed by either aldolase (e.g. FrbC)-catalyzed condensation to PnMM, aminotransferase (e.g. PalB)-catalyzed formation of PnAla, dehydrogenase (e.g. VlpB)-catalyzed reduction to PnLac, degradative reactions (PalA or PhnX, shown in grey), or Ppd-catalyzed decarboxylation to afford PnAA. **Right side:** Post-Ppd transformations include aldolase (e.g. RhiG), reductase (e.g. PhpC)-catalyzed formation of HEP, AEPT-catalyzed formation of AEP, and aldehyde reductase (e.g. PhnY)-catalyzed PnAc formation. Note that the PnAA-to-PnAc transformation can also be catalyzed by the α-ketoglutarate-dependent dioxygenase and PnhY* homolog FzmG. PFAM abbreviations and identification numbers of each enzyme are in square brackets. Dotted arrows represent reactions not biochemically characterized with purified enzyme. **Abbreviations:** PEP, phosphoenolpyruvate; PnPy, phosphonopyruvate; PnAA, phosphonoacetaldehyde; PnLac, phosphonolactate; PnMM, phosphonomethylmalate; PnAla, phosphonoalanine; PnAc, phosphonoacetate; AEP, 2-aminoethylphosphonate; HEP, 2-hydroxyethylphosphonate; KHPnPA, 2-keto-4-hydroxy-5-phosphonopentanoate; Ppm, PEP mutase; Ppd, PnPy decarboxylase; AEPT, AEP transaminase.

Since the 1959[8] discovery of AEP as a ubiquitous biological phosphonate and subsequent identification of its three-enzyme pathway (Ppm-Ppd-AEPT), several additional enzymatic branch points emanating from either PnPy or PnAA have been characterized. In addition to AEPT, three other post-Ppd enzymes have been biochemically verified (**Figure 1, right side**): (i) an aldolase RhiG in the rhizocticin biosynthetic pathway catalyzes decarboxylative aldol condensation of oxaloacetate and PnAA to afford 2-keto-4-hydroxy-5-phosphonopentanoate (KHPnPA);[9] (ii) Fe-containing alcohol dehydrogenases like PhpC reduce PnAA to 2-hydroxyethylphosphonate (HEP);[10] and (iii) aldehyde dehydrogenase-catalyzed oxidation of PnAA to phosphonoacetate (PnAc) has been biochemically characterized for PhnY of the phosphonoalanine (PnAla) degradation pathway,[11] but is also predicted for the biosynthesis of *O*-phosphonoacetic acid serine and some phosphonolipids.[2, 12] Although a PhnY-like reaction is also catalyzed by the α-ketoglutarate-dependent dioxygenase FzmG en route to fosfazinomycins, FzmG and homologs like PhnY* more commonly catalyze hydroxylation of a phosphonate α-carbon.[13, 14] Three enzymes are thought to act directly on PnPy and therefore do not require the action of Ppd (**Figure 1, left side**). Of these three, only the aldol-like condensation with acetyl-CoA to generate phosphonomethylmalate (PnMM) has been biochemically verified for the homocitrate synthase homologs FrbC and Pfs2.[15, 16] In contrast, aspartate aminotransferase homologs like PalB have yet to be characterized in vitro.[17, 18] Similarly, activity of the dehydrogenase VlpB has only been inferred from studies of the valinophos gene cluster in which phosphonolactate (PnLac) was isolated from spent media.[19] Although the ubiquitous and well-characterized enzymes PalA and PhnX can also act on PnPy and PnAA respectively, these enzymes are catabolic rather than biosynthetic. Overall, these seven biosynthetic branch points may be broadly classified as hydride transfer, amino transfer, or carbon-carbon bond formation.

For several decades genome mining has played an increasing role in metabolite discovery, with particular impact in microbial natural products.[20] New and more accessible tools like antiSMASH,[21] PRISM,[22] and BiG-SCAPE[23] are available for identifying and classifying biosynthetic gene clusters (BGCs). These tools are also more generally useful for classifying gene neighborhoods to glean insights from massive amounts of sequence data, as shown by the recent application of BiG-SCAPE to the computational characterization of Fe-S flavoenzymes.[24] The apparently near-universal origin of biological phosphonates from the action of the Ppm enzyme offers an opportunity for such classification. A landmark investigation of Ppm sequence data indicated that about 5% of bacterial genomes encode Ppm, and about half are likely to be involved in phosphonolipid of phosphonoglycan biosynthesis.[2] This study also revealed a strong correlation between gene neighborhood and Ppm sequence. Ppm sequence diversity has since been used to guide the discovery of several new phosphonates including valinophos and phosphonoalamides, illustrating the utility of genome mining for phosphonate discovery.[18, 19]

Despite the success of Ppm sequence-based discovery, several general features of early-stage phosphonate biosynthesis remain unclear. Specifically, the relative abundance of each biosynthetic branch point remains unknown. Moreover, *ppm*-containing gene neighborhoods that do not encode any of the seven branch points may represent fertile hunting grounds for new phosphonate chemistry. Here we employ computational tools to inventory these seven phosphonate biosynthetic branch points in *ppm*-containing gene neighborhoods from NCBI RefSeq complete bacterial genomes, as well as archaea and viruses. The most common predicted intermediates include AEP (56%), PnAla (9.2%), HEP (8.5%), and PnAc (6.2%). Significantly, ~13% of genomes examined encode none of these seven gateway enzymes and may generate novel phosphonates like phosphonofructose and phosphonoacrylate. Other surprises include the apparent biosynthetic use of C-P lyase genes and a virus-encoded AEP pathway.

## METHODS

### Identification of Ppm-encoding genes

Genomes were downloaded from the National Center for Biotechnology Information (NCBI) RefSeq database in June 2020, including complete bacteria (17,258), archaea (1,077), fungi (326), protists (97), plants (124), and animals (556) (**Table S1**). In addition we scanned 364 ‘huge phage’ genomes[25] and 501 draft contig metagenome-assembled genomes (MAGs) of Nucleo-Cytoplasmic Large DNA viruses (NCLDVs)[26]. Phosphoenolpyruvate mutase (Ppm) protein sequences were identified by hmmsearch (HMMER 3.1b2, February 2015; http://hmmer.org) for the Pfam family PEP_mutase (PF13714.7). Hits that aligned with at least 70% of profile HMMs were included. Sequences were then filtered for the presence of the Ppm-specific EDKX5NS motif [27] using the Perl script *ps_scan.pl* version 1.86 [28].

### Collection and network analysis of phosphonate biosynthetic gene neighborhoods

Protein accession numbers were used to extract the corresponding RefSeq genome assembly accession numbers from feature tables and then download the full RefSeq assembly files in Genbank format for each identified assembly. Genome coordinates were extracted from each feature table to include *ppm* and five flanking genes on each side, and corresponding nucleotide accession numbers were used to download genbank files via batch entrez. A Python script was used to take the identified coordinates and extract each gene neighborhood from full GenBank files and output a collection of gene neighborhood GenBank files as input for BiG-SCAPE.[23] The files used for BiG-SCAPE analysis comprised *ppm* gene neighborhoods of 869 complete bacterial genomes, 12 archaea, and 1 virus, for a total of 882 gene neighborhoods.

### Phylogenetic tree construction

The 424 bacterial Ppm protein sequences, the 12 Ppm sequences from the archaeal gene neighborhoods, and the single viral Ppm sequence (**Table S1**) were aligned against the PF13714 HMM using hmmalign. After removing unaligned and indel regions we were left with a 91 amino acid alignment of 432 Ppm sequences. A phylogenetic tree was constructed with FastTree [29] and the midpoint rooted tree was constructed using the phytools package in R [30]. The phylogenetic tree was visualized and annotated using the interactive Tree of Life (iTOL) [31].

### Classifying Ppm-encoding neighborhoods based on early biosynthetic branch point

The 882 gene neighborhoods were also classified according to their predicted early biosynthetic branch points following the Ppm-catalyzed formation of PnPy common to all neighborhoods. This classification was performed based on the presence of the key enzymes outlined in **Figure 1**, and PFAM domains for each were identified by hmmscan (HMMER 3.1b2, February 2015; http://hmmer.org). Three of these enzymes act on PnPy and therefore only need Ppm co-encoded in the gene neighborhood (PFAM in parentheses): aminotransferase PalB (Aminotran_1_2 PF00155.22) to afford PnAla[17, 18], phosphonomethylmalate synthase FrbC (HMGL-like PF00682.20)[15], and short-chain dehydrogenase VlpB (2-Hacid_dh_C PF02826.2) to generate PnLac[19]. The remaining four enzymes act after PnPy decarboxylase (Ppd, PF002775 or PF002776)[6]: AEP transaminase PhnW (Aminotran_5 PF00266.20)[32], aldolase RhiG (HMGL-like PF00682.20)[9], phosphonoacetaldehyde dehydrogenase PhnY (Aldedh PF00171.23)[11], and phosphonoacetaldehyde reductase PhpC (Fe-ADH PF00465.20)[33].

Each pathway was classified according to the presence and absence of encoded enzymes (in addition to Ppm) required for each of the seven metabolic outcomes shown in **Figure 1**. For example, a gene neighborhood was classified as ‘AEPT’ if it encoded (in addition to Ppm) Ppd and AEPT but did not encode the competing RhiG, PhpC, and PhnY homologs. A series of Python scripts were written to break each gene into individual protein sequences, use hmmscan to find all PFAM domains and assign them to each gene neighbourhood, identify the presence/absence of defined enzyme domains, and match each gene neighborhood to its corresponding BiG-SCAPE gene cluster family (GCF) number. The network was visualized in Cytoscape and manually curated for accurate branch point assignment of GCFs. The branch point inventory shown in Figure 2B was constructed by treating all BGCs possessing the same Ppm sequence as a single data point, such that a total of 434 rather than 822 BGCs are represented.

**Figure 2.**
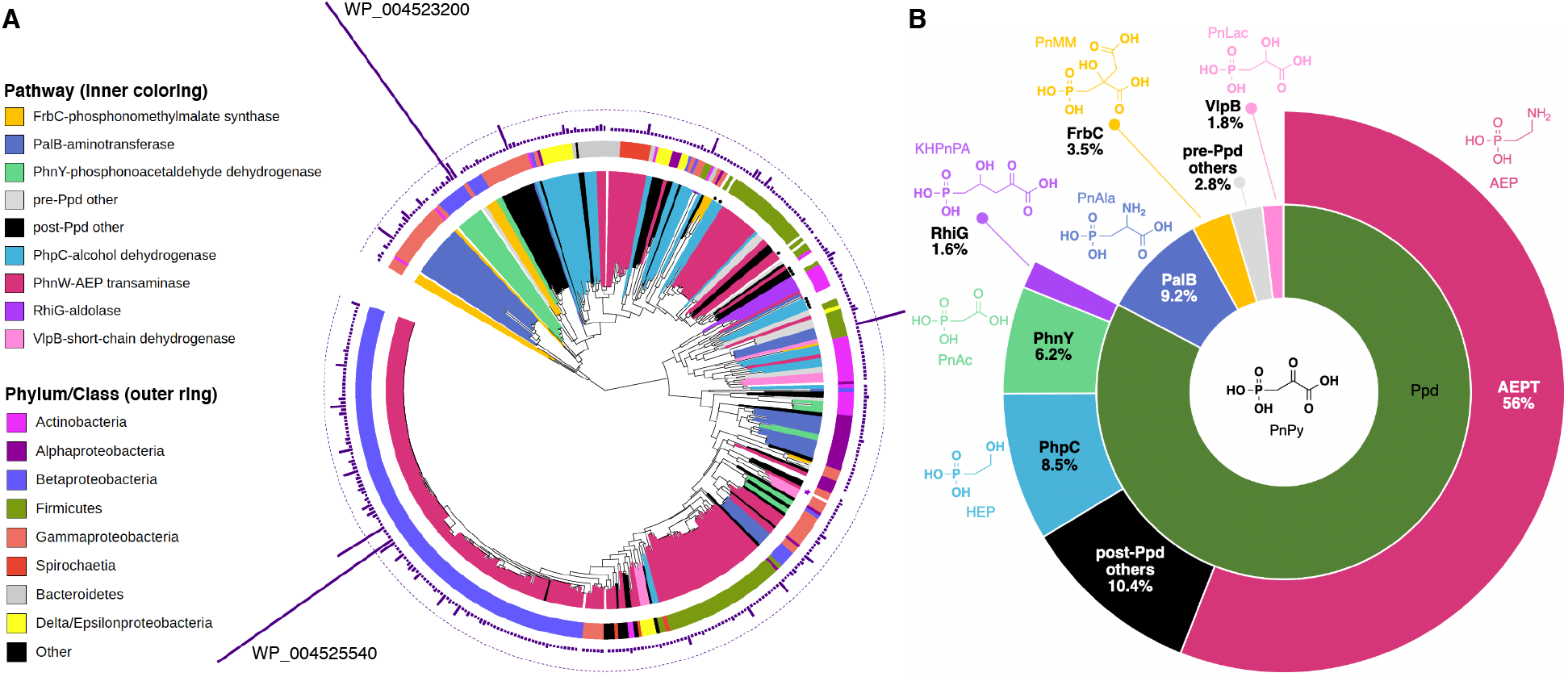
Classification of phosphonate biosynthetic genes. A) Phylogram of 424 unique Ppm sequences from RefSeq bacterial complete genomes, plus 12 archaeal (black dots) and a single viral Ppm sequence (purple star). Inner coloring represents the presence of neighboring genes encoding early-stage branch point transformations; the colors in the outer ring represent bacterial phylum or class. The outmost ring of bars represents the number of times a given sequence was found in RefSeq genomes, with the dotted line marking a count of 10. The two largest bars represent sequences WP_004523200 and WP_004525540. B) Sunburst chart of early branch point phosphonate biosynthetic steps based on 434 gene neighborhoods; BGCs possessing the same Ppm sequence (e.g. WP_004523200) are treated as a single data point. Starting from Ppm-generated PnPy, the inner ring illustrates that most PnPy (~83%, dark green ‘Ppd’) is likely transformed to PnAA based on the presence of a gene encoding Ppd. The outer ring illustrates that post-Ppd transformations are dominated by AEPT (red). The color scheme of B matches the inner ring of A.

## RESULTS AND DISCUSSION

### The microbial Ppm sequence inventory is dominated by *Burkholderia*

Consistent with previous work[2] we identified at least one Ppm-encoding gene in 4.2% of complete bacterial genomes, 1.6% of archaea, 12% of protists, and 3.8% of animals (**Table S1**). Although fungal phosphonates have been reported[34], we could not find Ppm sequences in fungal genomes, nor could we identify Ppm encoded by plant or viral genomes in the NCBI RefSeq database. Surprisingly, we detected Ppm encoded by one contig of a metagenomically-assembled genome of a Nucleo-Cytoplasmic Large DNA virus (NCLDV).[26] The vast majority of Ppm sequences (424 of 490) were bacterial, and many occurred in multiple genomes for a total of 869 Ppm-encoding gene neighborhoods in 723 bacterial genomes. Although most Ppm sequences were found in a single genome, several were found in multiple genomes. In particular, two Ppm sequences (WP_004523200 and WP_004525540) together accounted for over 200 gene neighborhoods in different genome assemblies of *Burkholderia pseudomallei*; however, the phylogenetic separation of these two sequences in distinct clades implies different phosphonate products (**Figure 2A**).[2] We also noted the previously-described abundance of Proteobacteria and in particular the genus *Burkholderia*, which comprised almost a quarter of the 424 bacterial Ppm sequences analyzed (**Figure 2A**).

### Gene neighborhoods predominantly encode transamination and hydride transfer

We classified 882 *ppm*-containing gene neighborhoods (*ppm* plus five genes on each side) according to the presence of the known early-stage phosphonate biosynthetic genes shown in **Figure 1**. After treating gene neighborhoods possessing the same Ppm sequence as a single data point (e.g. all 103 neighborhoods possessing the Ppm sequence WP_004523200 were counted as a single pathway leading to PnAc), we were left with 434 gene neighborhoods. The results of this classification are summarized in **Figure 2B**. As expected, *ppd* is found in the vast majority of the gene neighborhoods studied, indicating that ~83% of phosphonate biosynthetic pathways proceed through the intermediacy of PnAA. The next most common transformation of PnPy (in 9.2% of gene neighborhoods) is predicted to be catalyzed by PalB-like aminotransferases (PF00155 ‘Aminotran_1_2’) to afford PnAla. Although PalB belongs to a degradative pathway,[17] a family of homologous aminotransferases were recently implicated in phosphonoalamide biosynthesis.[18] The citrate synthase-like aldol reaction catalyzed by FrbC homologs as well as the VlpB-like reduction to PnLac together accounted for only 5.3% of the gene neighborhoods. Also as expected, the vast majority of post-Ppd reactions involve transamination catalyzed by AEPT, which is an aminotransferase (PF00266 ‘Aminotran_5’) distinct from PalB homologs. Overall, AEP is predicted to arise as an intermediate in more than half of all phosphonate biosynthetic pathways. The next most common known intermediate appears to be HEP, which accounts for 8.5% of neighborhoods. PhnY-generated PnAc was found in 6.2% of gene neighborhoods and putative RhiG-like aldol reactions in only 1.6%. The relative scarcity of HEP was surprising and may reflect an underrepresentation of marine sequences; HEP occurs in several natural product biosynthetic pathways (fosfomycin, phosphinothricin, dehydrophos, argolaphos), as a major polysaccharide component of marine dissolved organic matter [35], and a precursor to methylphosphonate en route to methane production in the ocean.[36] In summary, most gene neighborhoods encoded for early-stage amino (~65%) or hydride (~16%) transfer, with minor contributions from aldol-type carbon-carbon bond-forming reactions (~5%). The remaining ~13% of gene neighborhoods did not possess any of the known genes depicted in **Figure 1**, suggesting possible novel transformations and phosphonate products.

### Network analysis reveals several large gene cluster families

Although the above analysis classified key early branch points in phosphonate biosynthetic gene neighborhoods, it did not capture diversity within each group. For example, how varied are the gene neighborhoods comprising the 56% that appear capable of making AEP? To address this question we used BiG-SCAPE network analysis to classify the 882 gene neighborhoods into 85 different gene cluster families (GCFs, **Figure 3**).

**Figure 3.**
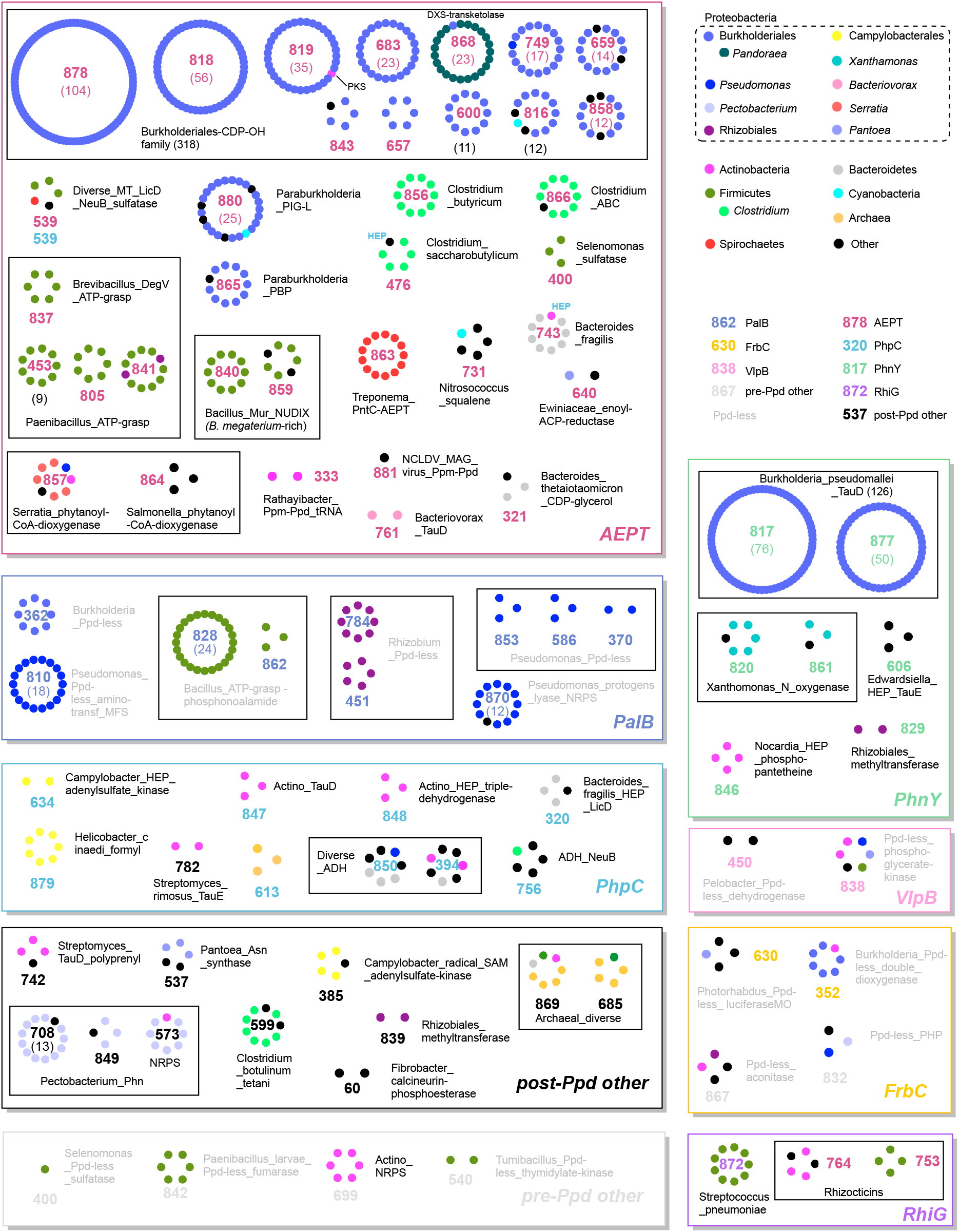
Network of 882 *ppm*-containing gene neighborhoods grouped by BiG-SCAPE into 85 gene cluster families (GCFs). Each dot represents a gene neighborhood colored according to organism type (e.g. phylum, genus). GCFs are organized into labeled boxes and colored by biosynthetic branch point. Each GCF is accompanied by a number (for identifying in BiG-SCAPE) and a descriptive name.

Half of the neighborhoods evaluated (444 of 882) belonged to AEPT- or PhnY-encoding *Burkholderia*-dominated gene neighborhoods, represented respectively by the ‘Burkholderiales_CDP-OH’ and ‘Burkholderia_TauD’ groups of GCFs (boxes in **Figure 3**). These larger groups respectively correspond to the Group 1 and 2 phosphonolipids described by Yu et al.[2] and are predicted to produce the 1-hydroxy substituted phosphonates shown in **Figure 4**. This predominance of *Burkholderia* may reflect database abundance due to medical importance rather than natural occurrence; for example, most of the 126 neighborhoods of GCFs 877 and 817 appear to be nearly identical strains of *Burkholderia pseudomallei*, the causative pathogen of meliodosis.[37] Furthermore, these two abundant groups contained the two Ppm sequences found in over 200 mostly *Burkholderia pseudomallei* gene neighborhoods (large bars in **Figure 2A**): WP_004523200 occurred in the majority of nearly identical PhnY-containing gene neighborhoods of GCFs 817 and 877 that also encoded a TauD homolog (**Figure 4 inset**); WP_004525540 occurred in the majority of near-identical AEPT-encoding neighborhoods of GCFs 878 and 818. The latter two neighborhoods belong to a larger ‘Burkholderiales_CDP-OH’ group of twelve GCFs totaling 318 (36%) slightly more diverse neighborhoods. As shown in **Figure 4**, this group has a common seven-gene core that encodes, in addition to the three AEP-producing enzymes Ppm, Ppd, and AEPT: (i) a putative α-ketoglutarate-dependent iron(II) oxygenase, (ii) two phosphonyl tailoring cytidylyltransferases (PntCs), one of which is fused to Ppm as often observed in phosphonate biosynthetic gene clusters,[38] and (iii) two genes associated with phospholipid biosynthesis. Interestingly, despite their prevalence and detection of *ppm* gene expression,[39] phosphonolipids have not been characterized from either of these large groups; however, 1-hydroxy-AEP is attached to a phosphonosphingolipid from *Bacteriovorax stolpii* (formerly *Bdellovibrio stolpii*), but the gene cluster belongs to the more distantly related AEPT-encoding GCF 761 (**Figures 3 and 5**).[40]

**Figure 4.**
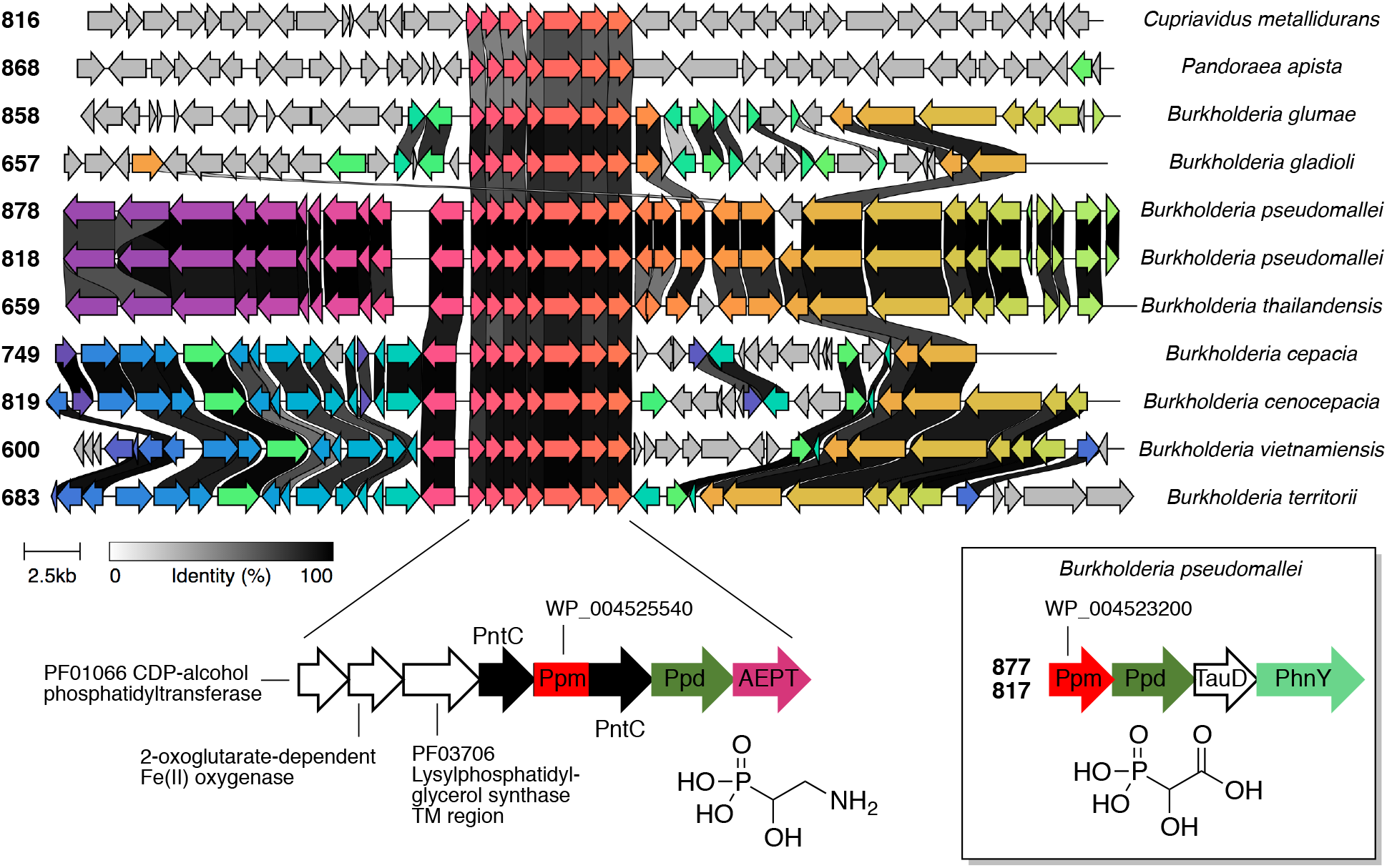
The two most abundant families of *ppm*-containing gene neighborhoods in the NCBI RefSeq database. Gene neighborhoods representative of the 318 ‘Burkholderiales-CDP-OH family’ members are shown aligned (with arbitrary color scheme) using clinker[50] to highlight the conserved seven-gene core, which is expanded below to show annotations and putative phosphonate product. The accession numbers WP_004525540 and WP_004523200 represent the most common Ppm in AEPT- and PhnY-containing clusters, respectively. Gene cluster family numbers corresponding to those in Figure 3 are provided to the left of each neighborhood. Inset illustrates the four-gene core representative of gene cluster families 877 and 817 with the putative phosphonate product. NCBI RefSeq accession numbers along with the corresponding Ppm protein accession numbers for each gene neighborhood are as follows: *Cupriavidus metallidurans* strain FDAARGOS_675, NZ_CP046331, WP_011516537; *Pandoraea apista* strain AU2161, NZ_CP011501, WP_042112012; *Burkholderia glumae* strain 257sh-1 chromosome 2, NZ_CP035901, WP_017423874; *Burkholderia gladioli* strain ATCC 10248 chromosome 2, NZ_CP009322, WP_036031650; *Burkholderia pseudomallei* isolate UKMPMC2000 chromosome 2, NZ_LR595895, WP_004525540; *Burkholderia pseudomallei* B03 chromosome 2, NZ_CP009150, WP_004525540; *Burkholderia thailandensis* E254 chromosome 2, NZ_CP004382, WP_043296855; *Burkholderia cepacia* ATCC 25416 chromosome 2, NZ_CP007748, WP_021162084; *Burkholderia cenocepacia* strain FL-5-3-30-S1-D7 chromosome 2, NZ_CP013396, WP_023477253; *Burkholderia vietnamiensis* strain AU1233 chromosome 1, NZ_CP013433, WP_060044431; *Burkholderia territorii* strain RF8-non-BP5 chromosome 2, NZ_CP013365, WP_059507240.

**Figure 5.**
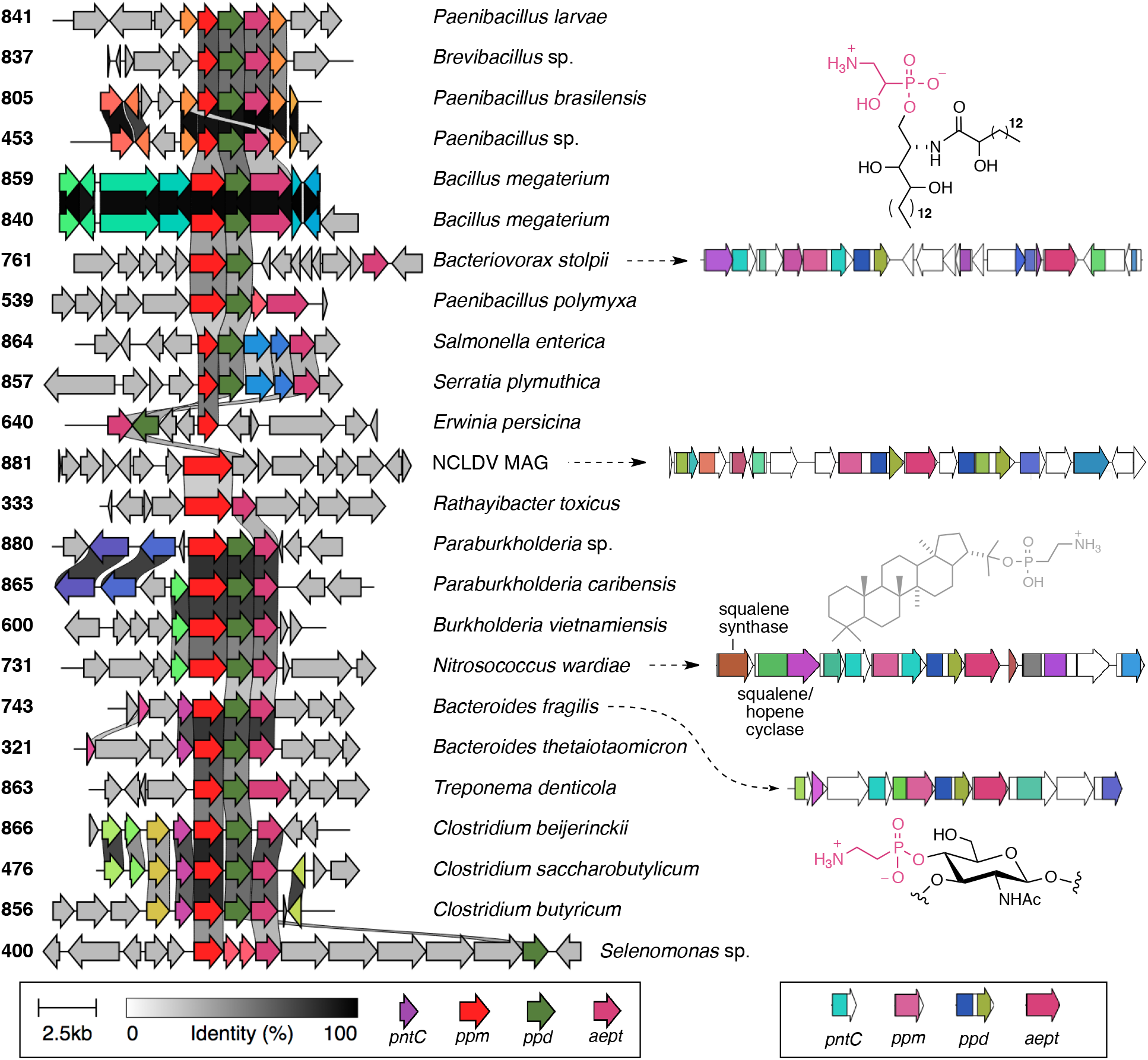
Diversity of AEPT-encoding gene neighborhoods. Representative neighborhoods for each family are shown on the left aligned by clinker[41] to highlight the key three-gene subcluster encoding AEP. On the right side are selected neighborhoods that include individual Pfam domains as colored by BiG-SCAPE and known (black) or proposed (gray) phosphonate products. NCBI RefSeq accession numbers for each family along with the corresponding Ppm protein accession numbers for each gene neighborhood are as follows: 841, *Paenibacillus larvae* subsp. larvae strain Eric_III, NZ_CP019655, WP_023485328; 837, *Brevibacillus* sp. 7WMA2, NZ_CP048799, WP_013336797; 805 *Paenibacillus brasilensis* strain KACC 13842, NZ_CP09363115, WP_025716510; 453, *Paenibacillus* sp. IHB B3084, NZ_CP013203, WP_134911858; 859, *Bacillus megaterium* strain FDU301, NZ_CP045272, WP_171777684; 840, *Bacillus megaterium* strain **S188**, NZ_CP049296, WP_164797121; 761, *Bacteriovorax stolpii* strain DSM 12778, NZ_CP025704, WP_102244880; 539, *Paenibacillus polymyxa* E681, NZ_014483, WP_013312267; 864, *Salmonella enterica* strain 85-0120, NZ_CP054715, WP_171775701; 857, *Serratia plymuthica* S13, NZ_021659, WP_004946681; 640, *Erwinia persicina* strain B64, NZ_CP022725, WP_118663913; 333, *Rathayibacter toxicus* strain WAC3373, NZ_CP013292, WP_052493485; 880, *Paraburkholderia* sp. Msb3 isolate PDMSB31, NZ_LR699554, WP_165187990; 865, *Paraburkholderia carabensis* strain 852011, NZ_CP015959, WP_062917090; 600, *Burkholderia vietnamiensis* strain AU1233, NZ_CP013433, WP_060044431; 731, *Nitrosococcus wardiae* strain D1FHS, NZ_CP038033, WP_134359243; 743, *Bacteroides fragilis* strain NCTC 9343, NC_003228, WP_011202610; 321, *Bacteroides thetaiotaomicron* strain 7330, NZ_CP012937, WP_008767814; 863, *Treponema denticola* ATCC 35405, NZ_002967, WP_002679012; 866, *Clostridium beijerinckii* isolate C. beijerinckii DSM 6423, NZ_LN908213, WP_077842376; 476, *Clostridium saccharobutylicum* strain NCP 195, NZ_CP016092, WP_022744173; 856, *Clostridium butyricum* strain 4-1, NZ_CP039705, WP_002579294; 400, *Selenomonas* sp. oral taxon 478 strain F0592, NZ_CP012071, WP_050342316.

### The large group of AEPT-encoding gene neighborhoods is diverse

Apart from the Group 1 ‘Burkholderiales_CDP-OH’ phosphonolipids, the AEPT-containing gene neighborhoods are diverse (**Figure 5**). These neighborhoods include those encoding two structurally characterized phosphonates: the above-mentioned *B. stolpii* phosphonosphingolipid [40] and the phosphonoglycan component of the polysaccharide B virulence factor from *Bacteroides fragilis*.[41] Also included are the *Selenomonas* species of GCF 400 that encode multiple sulfatases and correspond to the previously assigned Group 4 phosphonolipids.[2] As evident from the similarity of aligned genes in **Figure 5**, several gene cluster families are related. For example, GCFs 841, 837, 805, and 453 are all Paenibacillaceae encoding two ATP-grasp amino acid ligases flanking the AEP biosynthetic core and therefore may produce a peptide. The ATP-grasp and Mur ligase domains of GCFs 859 and 840 suggest phosphonylated peptidoglycan. Although GCFs 864 and 857, exemplified by the human pathogen *Salmonella enterica*, possess the classical three-gene AEP core, they also encode an aminotransferase (PF00155) associated with PnAla production.[18] Squalene synthase and cyclases encoded by *Nitrosococcus* species of GCF 731 hint at a possible phosphonoterpene product as previously speculated.[42] This comparison of AEPT-encoding gene clusters also showcases diverse gene fusions. For example, Ppm can be encoded alone (GCFs 764 and 753) or as a fusion to a PntC cytidylyltransferase (e.g. PF12804 in GCF 761, or PF01467 in GCF 743) or Ppd (GCFs 333 and 881). Curiously, GCF 881 is a single gene neighborhood from a metagenomically assembled genome of a Nucleo-Cytoplasmic Large DNA Virus (NCLDV).[26] Immediately downstream of the AEP core are genes implicated in aminosugar synthesis consistent with production of a viral phosphonoglycan.

### The unexplored diversity of PhnY-, PalB-, and PhpC-encoding pathways

Phosphonoacetate (PnAc) is an intermediate in only two characterized biosynthetic pathways: the fosfazinomycins [13] and *O*-phosphonoacetic acid serine[12]. In fosfazinomycin biosynthesis the transformation of PnAA to PnAc is probably due to the activity of FzmG, an a-ketoglutarate Fe(II)-dependent oxygenase whose primary role in this pathway is hydroxylation at C-1 of methylated PnAc.[13] The more common route to PnAc is via PhnY-like aldehyde dehydrogenases that have been characterized from phosphonate degradation pathways.[11] Evidence for biosynthetic roles for PhnY-like enzymes has been found in: (i) the PhpJ transformation of phosphonoformaldehyde to phosphonoformate in phosphintothricin biosynthesis[33], and (ii) PnAc formation in *O*-phosphonoacetic acid serine biosynthesis in *Streptomyces* sp. NRRL F-525.[12] The latter gene neighborhood is shown in **Figure S1A** along with other GCFs encoding PhnY-like aldehyde dehydrogenases, highlighting diversity and some interesting features. For example, the actinomycete-dominated GCF 846 and *Bradyrhizobium* species in GCF 829 encode non-ribosomal peptide synthetase (NRPS) systems within 15 genes of *ppm*, and the former encodes putative adenylation and thiolation domains within two genes of the encoded aldehyde dehydrogenase. Plant pathogens like *Xanthomonas vasicola* dominate GCFs 820 and 861, each of which possess a seven-gene core encoding an AurF-like *N*-oxygenase (PF11583), glutamine amidotransferase and asparagine synthase.

PalB-encoding families also very diverse and include GCF 870 of *Pseudomonas protegens* strains encoding NRPS systems within two genes of *ppm* (**Figure S1B**). Notably, none of these families represent the previously discovered group of phosphonalamides produced by various *Streptomyces* species and isolated from *Streptomyces* sp. NRRL B-2790, which are the only known phosphonate biosynthetic products of PalB-like activity.[18] The absence of the known phosphonoalamide gene clusters highlights the limitations of the current study using only bacterial complete genomes.

The formation of HEP catalyzed by PhpC-like iron-containing alcohol dehydrogenases appears in diverse families of gene neighborhoods that include pathogenic bacteria like *Helicobacter cinaedi* and *Campylobacter jejuni* (**Figure S1C**). Although this transformation is important in several *Streptomyces* natural products like fosfomycin, phosphinothricin, dehydrophos, and argolaphos,[1] none of the encoding gene clusters were found in the NCBI complete genome database. Of the ten gene cluster families identified, only two have yielded isolated phosphonate products: the methylphosphonate adduct from the marine archaeon *Nitrosopumilus maritimus*,[36] and the HEP conjugate purified from exopolysaccharide of *Stackebrandtia nassauensis*.[43] Interestingly, the two *Streptomyces rimosus* strains comprising GCF 782 encode not only a PhpC homolog, but also VlpB and PhnY homologs. This is one example highlighting the ambiguities resulting from our classification system because this cluster could be attributed to any number of groups and therefore can benefit from manual curation. Specifically, the presence of a possible 2-hydroxyethylphosphonate dioxygenase (HEPD) homolog of PhpD implies intermediacy of HEP in a manner reminiscent of phosphinothricin biosynthesis in *Streptomyces viridochromogenes*. In this case HEP would be oxidized by PhpC to phosphonoformaldehyde followed by PhnY-catalyzed formation of phosphonoformate analogous to the proposed action of PhpJ.[33]

### VlpB-like dehyrogenases: beyond valinophos

To date PnLac has only been isolated in spent media of the valinophos producer *Streptomyces durhamensis* B-3309, and the dehydrogenase VlpB was proposed to catalyze its formation from PnPy.[19] Very similar gene clusters are found in two other *Streptomyces* strains of GCF 838: *S. atratus* strain SCSIO_ZH16 and *Streptomyces* sp. WAC 01529. Both of these clusters possess the *vlpABCDE* core proposed to generate 2,3-dihydroxypropylphosphonic acid (DHPPA) (**Figure S2**). Three other members of this family, all Proteobacteria, expand upon the five-gene valinophos core. The lone Firmicutes, *Tumebacillus algifaecus*, possesses a unique gene cluster that may encode the novel compound phosphonofructose, an analog of the sulfoglycolytic intermediate sulfofructose (**Figure S3**).[44]

### The aldolases: FrbC- and RhiG-encoding pathways

The GCFs encoding Ppm, Ppd, and FrbC en route to the intermediate PnMM diverge from the standard FR-900098-like pathway logic of GCF 867 (**Figure 6A**) to include more unusual gene combinations in GCFs 352 and 630 (**Figure S4**). Like the FR-900098 gene cluster, GCF 867 encodes aconitase and isocitrate dehydrogenase homologs en route to the α-ketoglutarate analog 2-oxo-4-phosphonobutyrate.[15] Most GCF 867 gene clusters do not encode an aminotransferase, but like FR-900098 biosynthesis may nonetheless generate the glutamate analog 2-amino-4-phosphonobutyrate via the action of a promiscuous enzyme encoded elsewhere in the genome. However, we identified a plasmid of *Agrobacterium tumefaciens* strain CFBP6625 that also encodes a PalB-like aminotransferase and therefore may facilitate transamination to 2-amino-4-phosphonobutyrate (**Figure 6A**). The *Burkholderia*-dominated GCF 352 is the largest of the four FrbC GCFs and possesses an unusual gene combination including two dioxygenases, lumazine synthase, and a member of the beta-keto acid cleavage enzyme (BKACE) family.[45] The *Photorhabdus* GCF 630 neighborhoods encode aconitase and luciferase-like monooxygenase, an FMN-dependent enzyme found in natural product biosynthetic pathways such as mensacarcin and neoabyssomicin.[46, 47]

**Figure 6.**
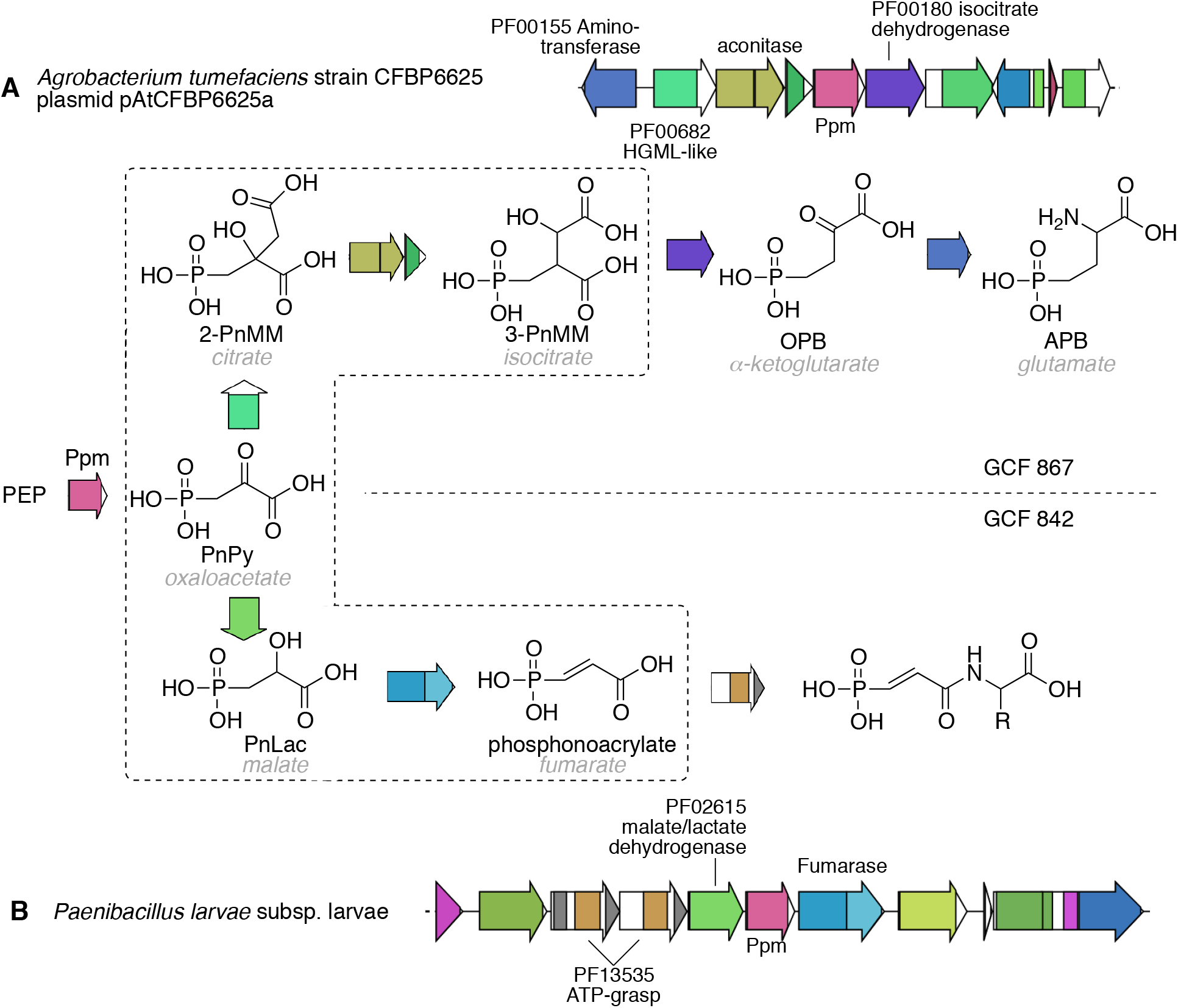
Possible citric acid cycle homologs in **A)** FrbC-like pathway of GCF 867 and **B)** unclassified GCF 842. The transformations mimicking those of the citric acid cycle are in the dotted box region, and the names of non-phosphonate analogs are italicized in grey. Abbreviations: 2-PnMM (2-phosphonomethylmalate), 3-PnMM (3-phosphonomethylmalate), OPB (2-oxo-4-phosphonobutyrate), APB (2-amino-4-phosphonobutyrate), PnLac (phosphonolactate), PnPy (phosphonopyruvate).

GCFs 764 and 753 (**Figure S4**) are very similar and were assigned as RhiG aldolase groups due to the tight clustering of genes in the order *ppd-rhiG-ppm* and the inclusion of the known rhizocticins producer *Bacillus subtilis* ATCC6633 in this group.[9] One distinguishing feature of these rhizocticin-like gene clusters is the apparent separation of Ppd domains (PF2775 and PF2776) onto individual proteins, which is occasionally observed in other gene clusters such as the *O*-phosphonoacetic acid serine cluster from PnAc-producing *Streptomyces* sp. strain NRRL F-525 (**Figure S1A**) [12]. The presence of an HMGL-like domain in the *Streptococcus pneumoniae* clusters of GCF 872 led to assignment as possible RhiG-encoding pathways, with an unusual tRNA synthetase and phosphopantetheine attachment site suggesting a peptidyl product (**Figure S5**). Notably this neighborhood is conserved across only ten of 81 complete NCBI RefSeq *S. pneumoniae* genomes, and the flanking recombinase/relaxase genes define potential gene cluster boundaries.

### Discovery potential in unclassified neighborhoods

Fifteen gene cluster families could not be assigned because they did not encode any of the seven branch point enzymes acting on either PnPy or PnAA shown in **Figure 1**. Most of these GCFs encoded Ppd (‘post-Ppd other’ in **Figure 3**) but several did not (‘pre-Ppd other’ in **Figure 3**). Surveying these unclassified families revealed several unusual features. For example, GCF 842 neighborhoods of *Paenibacillus larvae* encode Ppm flanked by malate dehydrogenase and fumarase (**Figure 6B**). Because PnPy is an analog of the citric acid cycle intermediate oxaloacetate, we predict that malate dehydrogenase will catalyze a VlpB-like reduction to PnLac followed by fumarase-catalyzed dehydration to the fumarate analog phosphonoacrylate. The adjacent ATP-grasp domains imply peptide bond formation with an amino acid as seen for the phosphonoalamides.[18] Interestingly, directing PnPy in the fumarase direction of the TCA cycle would contrast with the citrate synthase direction observed in FR-900098 biosynthesis and in FrbC-encoding families such as GCF 867 (**Figure 6**). GCF 540 encodes Ppm that is unusually flanked by thymidylate kinase and an uncharacterized nucleotidyltransferase (PF14907), while actinomycete-dominated GCF 699 has several gene neighborhoods encoding a PLP-dependent enzyme (PF00291) that may also catalyze a PalB-like transamination to PnAla (**Figure S4**).

Most of the uncharacterized diversity occurs in gene neighborhoods that encode Ppm and Ppd and therefore likely proceed through the intermediacy of PnAA (**Figure S4**). This includes the unusual occurrence of the C-P lyase *phn* operon[48]: GCFs 573, 708, and 849 encode Ppm, Ppd, and a possible dehydrogenase followed on the same strand by PhnGHIJKLMNP (**Figure S6**). The absence of both the PhnCDE transporter and the PhnO *N*-acetylation enzyme imply a function other than AEP import and degradation. The presence of an NAD(P)-dependent oxidoreductase after Ppm-Ppd implies HEP or PnAc formation using atypical enzymes. Several gene clusters in GCF 537 encode a hotdog-fold enzyme associated with A-factor biosynthesis, a phosphopantetheine attachment site, and asparagine synthase. Neighborhoods dominated by *Clostridium botulinum* and *C. tetani* in GCF 599 encode a methyltransferase and sugar biosynthetic enzymes. In *Campylobacter* strains of GCF 385, the *ppm-ppd* core is flanked by genes predicted to encode radical SAM and adenylsulfate kinase enzymes. *Fibrobacter succinogenes* of GCF 60 is known to produce an exopolysaccharide possessing a novel *N*-hydroxyethyl derivative of AEP.[49] However, the absence of AEPT suggests that AEP is not an intermediate but rather favors the reductive amination of PnPy with ethanolamine. The *Streptomyces* of GCF 742 possess a unique combination of polyprenyl synthetase, cytochrome P450, methyltransferase, and a TauD-like oxygenase. Archaea of GCFs 869 and 685 have diverse genes surrounding *ppm* and *ppd*, including calcineurin-like phosphoesterase (PF12850) and GDP-mannose 4,6-dehydratase (PF16383) domains in common with *Fibrobacter succinogenes*.

### Genomes with multiple Ppm-encoding neighborhoods

The vast majority of the 726 *ppm*-containing bacterial genomes evaluated possess either one (~80% of genomes) or two (~18% of genomes) *ppm*-containing gene neighborhoods, but several maintained three or four (SI file SI_multiply_ppm.xls), as previously noted[2]. Apart from a single *Streptomyces* genome (*S. griseochromogenes* ATCC 14511) with three neighborhoods, all genomes encoding more than two Ppm were *Burkholderia*, and four of the five genomes with four *ppm* genes were strains of *B. oklahomensis*. An analysis of these *Burkholderia* Ppm gene neighborhoods revealed 151 genomes with a single *ppm;* 119, 3, and 5 genomes housed two, three, and four *ppm*, respectively (**Figure S7**). All genomes encoded AEPT-containing GCFs belonging to the large ‘Burkholderiales CDP-OH family’ dominated by GCFs 818 and 878, but a clear combinatorial progression emerges with additional clusters. Specifically, the second cluster is always PhnY-containing GCFs 877 or 817, the third cluster belongs to PalB-like GCF 362, and the fourth cluster is GCF 352 that we assigned as FrbC-like (**Figure S7**). The significance of this expanded phosphonate biosynthetic capacity in *Burkholderia* is unclear, but the consistent accumulation of the same four classes of phosphonate biosynthetic gene cluster is intriguing.

### Conclusion

Phosphonates are an important but niche class of natural products and increasingly appreciated as important cell surface modifications in the form of phosphonolipids and phosphonoglycans. Because most biological phosphonates originate via Ppm catalysis, and Ppm sequence phylogeny correlates with biosynthetic product[2], discovery of new phosphonates has largely focused on *ppm*-guided actinomycete strain prioritization[19]. However, less is known about the diversity of *ppm* gene neighborhoods and how they might be classified in order to identify and prioritize overlooked gene clusters. Specifically, little is understood about the relative abundance of different branch points in the early steps of phosphonate biosynthesis. This study provides an inventory of these early branch points and highlights potential avenues of discovery. These highlights include: (i) a possible phosphonoacrylate intermediate, (ii) integration of the degradative C-P lyase (*phn* operon) components into biosynthetic gene clusters, (iii) a possible phosphonofructose biosynthetic pathway, and (iv) a virus-encoded phosphonate, which hints at the broader possibility of virus-encoded natural products. In summary, these results help contextualize current knowledge of early branch points of phosphonate biosynthesis and provide a framework to generate hypotheses and triage discovery efforts.

## Acknowledgements

We thank Monica Papinski for providing Python functions that we modified and incorporated into final scripts.

