## Supplemental Information for "An inventory of early branch points in microbial phosphonate biosynthesis"

**Table S1**. Identified Ppm-encoding genes and gene neighborhoods.

| RefSeq Assembly Type | Total RefSeq assemblies searched (.faa) | Unique Pfam hits from hmm-search | Unique Ppm hits after filtering for $EDKX_5NS$ motif | Number of assemblies encoding at least one Ppm (% total) | Number of clusters |
|---|---|---|---|---|---|
| Bacteria complete | 17,258 | 9693 | 424 | 723 (4.19%) | 869 |
| Virus (NCBI) | 10001 | 0 | | | |
| Huge_phage | 364 | 0 | | | |
| NCLDV_MAGs | 501 | 1 | 1 | 1 | 1 |
| Archaea | 1077 | 408 | 16 | 17 (1.6%) | 12 |
| Protists | 97 | 87 | 15 | 12 (12%) | |
| Fungi | 326 | 1370 | 0 | | |
| Plants | 124 | 577 | 0 | | |
| Animals | 556 | 85 | 34 | 21 (3.8%) | |

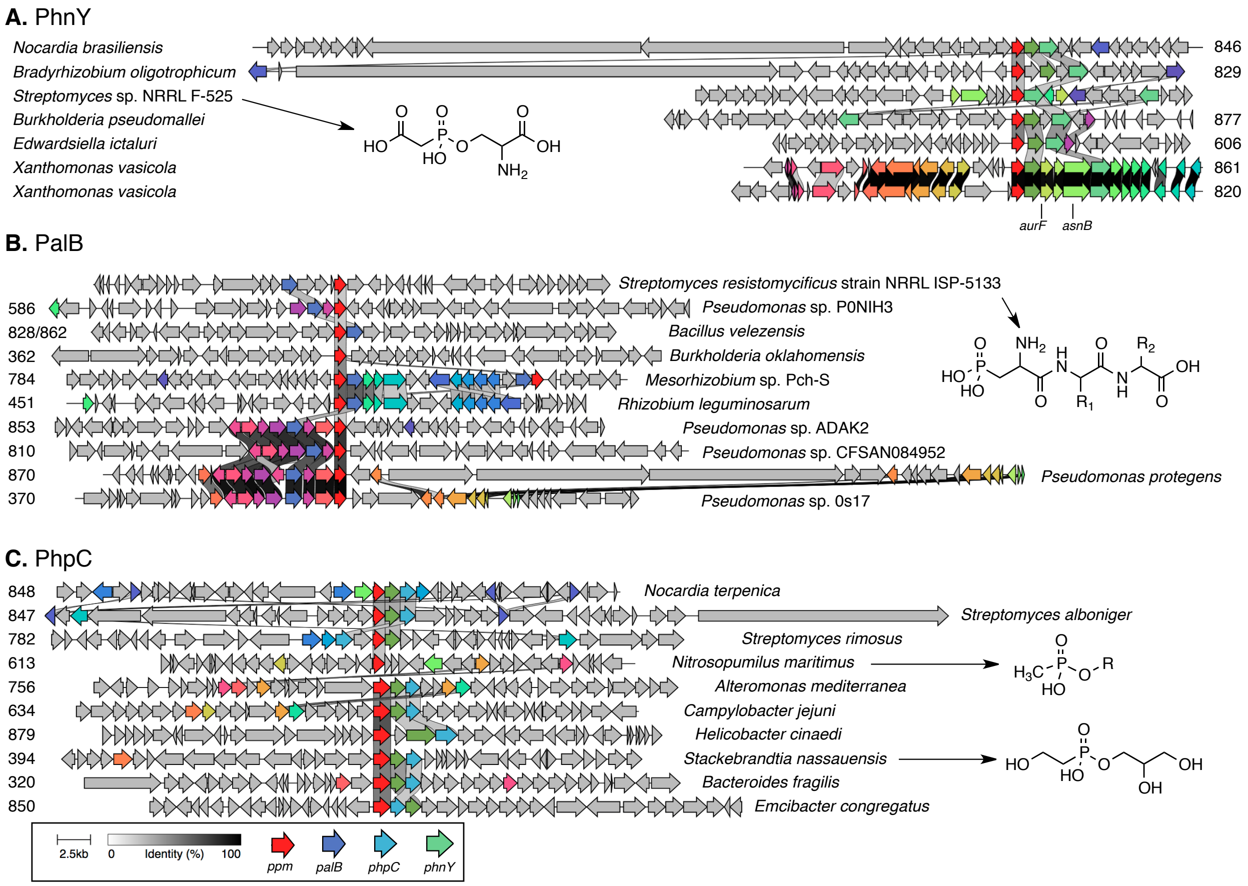


**Figure S1.** Summary of gene neighborhoods containing PhnY, PalB, or PhpC. A representative gene neighborhood for each gene cluster family is shown aligned with other family representatives and identity scoring performed using clinker. The detailed strains, nucleotide accession number, and Ppm accession number are as follows: **PhnY.** 606 - *Edwardsiella ictaluri* 93-146, NC_012779, WP_015871974; 820 - *Xanthomonas vasicola* pv. arecae strain NCPPB 2649, NZ_CP034653, WP_017116614; 829 - *Bradyrhizobium oligotrophicum* S58, NC_020453, WP_015664097; 846 - *Nocardia brasiliensis* ATCC 700358, NC_018681, WP_014985093; 861 - *Xanthomonas vasicola* strain NCPPB 1060, NZ_CP034649, WP_039437895; 877 - *Burkholderia pseudomallei* isolate UKMR15 chromosome 2, NZ_LR595899, WP_004523200; not included in network: *Streptomyces* sp. NRRL F-525, NZ_JNXE01000019, WP_051801207. **PalB.** *Streptomyces resistomycificus* strain NRRL ISP-5133, NZ_KL575589, WP_030038712; 362 - *Burkholderia oklahomensis* strain E0147, NZ_CP008726, WP_038801024; 370 - *Pseudomonas* sp. 0s17*,* NZ_AP014627, WP_060838450; 451 - *Rhizobium leguminosarum* strain Vaf-108 plasmid, NZ_CP018229, WP_062940190; 586 - *Pseudomonas* sp. P0NIH3, NZ_CP026386, WP_023629505; 784 - *Mesorhizobium* sp. Pch-S, NZ_CP029562, WP_129413975; 810 - *Pseudomonas* sp. CFSAN084952, NZ_CP045767, WP_032904173; 828/862 - *Bacillus velezensis* strain SRCM102755, NZ_CP028204, WP_014417091; 853 - *Pseudomonas* sp. ADAK2, NZ_CP052862, WP_169374082; 870 - *Pseudomonas protegens* Cab57, NZ_AP014522, WP_011060442. **PhpC**. 320 - *Bacteroides fragilis* strain DCMSKEJBY0001B, NZ_CP036546, WP_032537204; 394 - *Stackebrandtia nassauensis* DSM 44728, NC_013947, WP_041627074; 613 - *Nitrosopumilus maritimus* SCM1, NC_010085, WP_012214543; 634 - *Campylobacter jejuni* strain CJ031CC45, NZ_CP012211, WP_032603622; 756 - *Alteromonas mediterranea* strain PT15, NZ_CP041170, WP_141153524; 782 - *Streptomyces rimosus* strain ATCC 10970, NZ_CP023688, WP_030177674; 847 - *Streptomyces alboniger* strain ATCC 12461, NZ_CP023695, WP_070321175; 848 - *Nocardia terpenica* strain NC_YFY_NT001; NZ_CP023778, WP_098695139; 850 - *Emcibacter congregatus* strain ZYLT, NZ_CP041025, WP_099471844; 879 - *Helicobacter cinaedi* isolate MGYG-HGUT-01432, NZ_LR698961, WP_002956559.

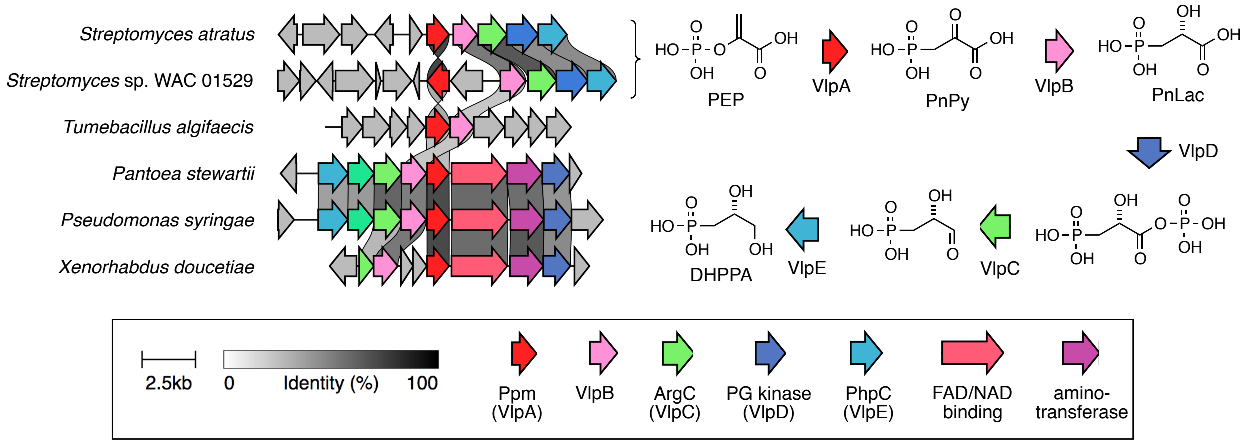


**Figure S2.** Gene neighborhoods comprising VlpB-encoding gene cluster family 838. At right are the first five steps of the valinophos biosynthetic pathway encoded by the two *Streptomyces* species. The nucleotide and Ppm accession numbers are as follows: *Streptomyces atratus* strain SCSI0_ZH16, NZ_CP027306, WP_114243022; *Streptomyces* sp. WAC 01549, NZ_CP029617, WP_125516761; *Tumebacillus algifaecus* strain THMBR28, NZ_CP022657, WP_094238279; *Pantoea stewartii* strain ZJ-FGZX1, NZ_CP049115, WP_133745451; *Pseudomonas syringae* CC1557, NZ_CP007014, WP_038401180; *Xenorhabdus doucetiae* strain FRM16, NZ_FO704550, WP_045973335.

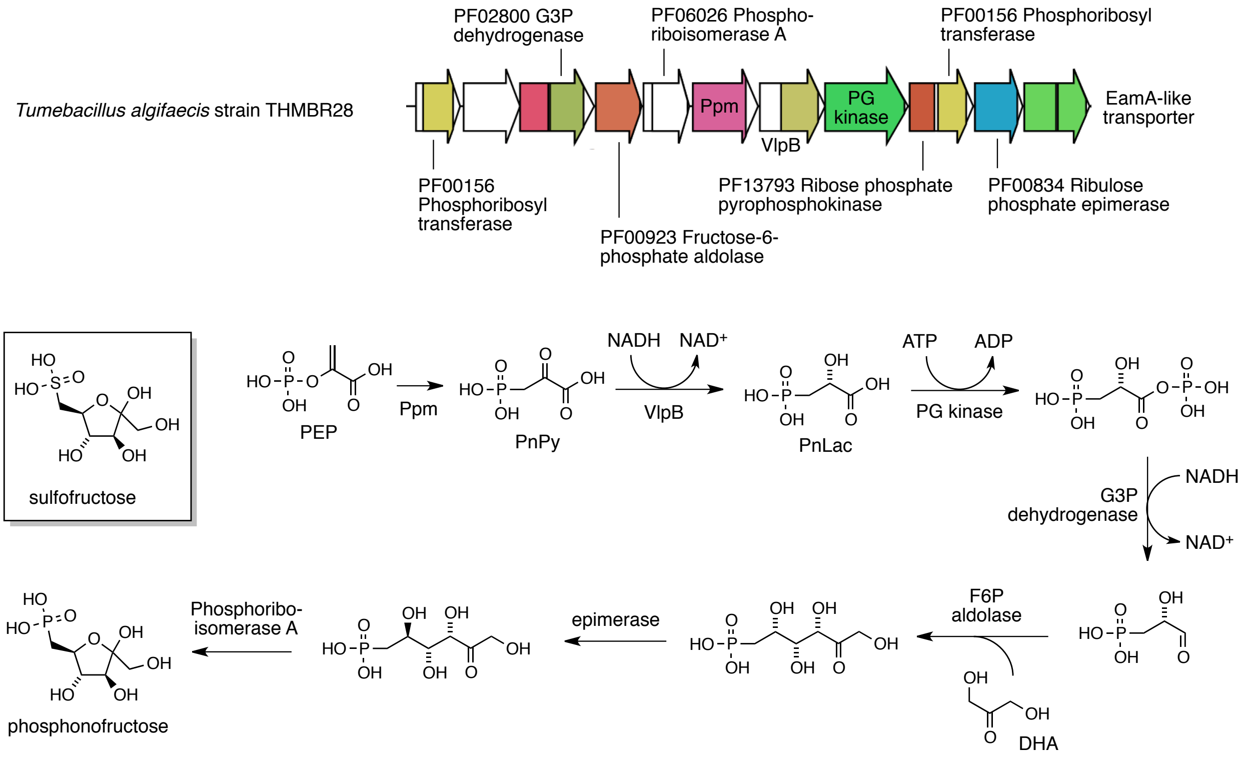


**Figure S3**. Gene neighborhood from *Tumebacillus algifaecis* (accession NZ_CP022657) in gene cluster family 838 and the proposed phosphonofructose biosynthetic pathway, illustrating comparison to the known sulfoglycolytic compound sulfofructose.

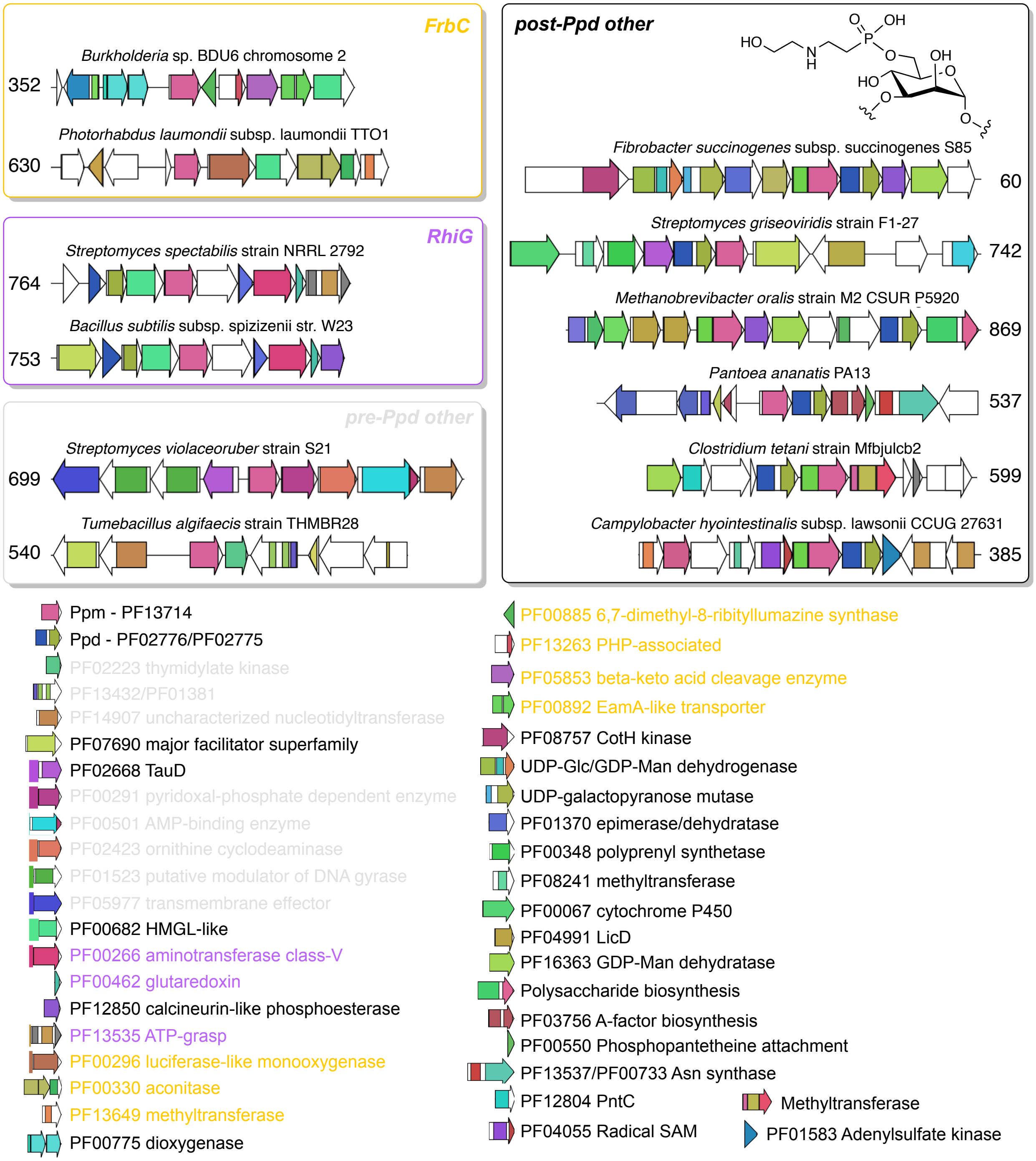


**Figure S4**. Representative *ppm*-containing gene neighborhoods from diverse GCFs. The phosphonate polysacharride isolated from *Fibrobacter succinogenes* is shown, although it has not been definitively linked to the gene cluster. Nucleotide and Ppm protein accession numbers are: **352 -** *Burkholderia* sp. BDU6, NZ_CP013387, WP_059472091; **630** - *Photorhabdus laumondii* subsp. laumondii TTO1, NC_005126, WP_011146132; **764** - S*treptomyces spectabilis* strain NRRL 2792, NZ_CP040916, WP_144322200; **753** - *Bacillus subtilis* subsp. spizizenii str. W23, NC_014479, WP_003223692; **699** - *Streptomyces violaceoruber* strain S21, NZ_CP020570, WP_030116101; **540** - *Tumebacillus algifaecis* strain THMBR28, NZ_CP022657, WP_094236718; **537** - *Pantoea ananatis* PA13, NC_017554, WP_014606798; **599** - *Clostridium tetani* strain Mfbjulcb2, NZ_CP027782, WP_115605286; **385** - *Campylobacter hyointestinalis*

subsp. lawsonii CCUG 27631, NZ_CP015576, WP_063997597; **742** - *Streptomyces griseoviridis* strain F1-27, NZ_CP034687, WP_127176566; **869** - *Methanobrevibacter oralis* strain M2 CSUR P5920, NZ_LT985106, WP_063720063; **60** - *Fibrobacter succinogenes* subsp. succinogenes S85NC_017448, WP_014545731.

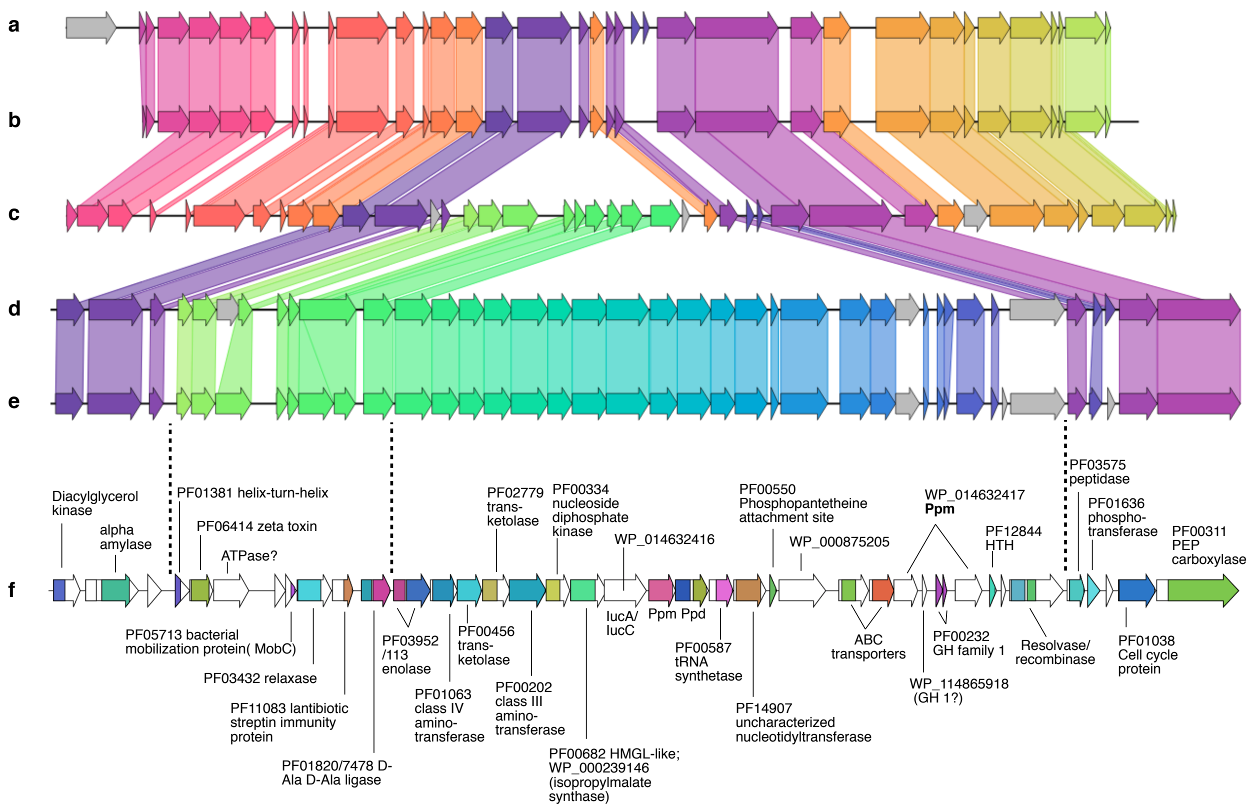


**Figure S5**. Comparison of GCF 872 gene neighborhoods in the following strains of *Streptococcus pneumoniae* (accession numbers in parentheses) **a.** Hu15 (CP020551), b. Hungary19A-6 (CP000936), c. 2245STDY5775603 (LR216036), d. M26368 (CP031246), e. OXC141 (NC_017592), f. OXC141 showing annotations and coloring from BiG-SCAPE. Possible phosphonate biosynthetic gene cluster boundaries are denoted with dotted lines.

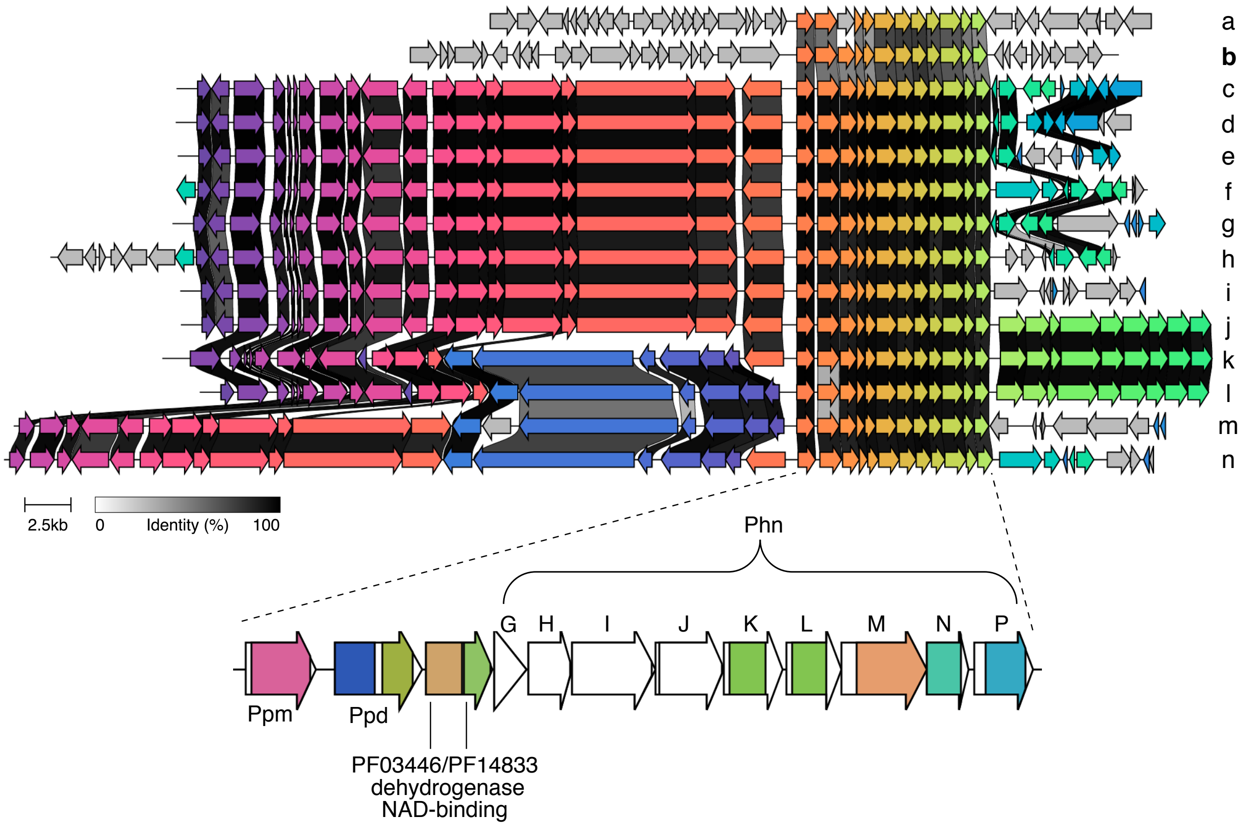


**Figure S6.** Selected strains that possess the *phnGHIJKLMNP* operon adjacent to *ppm-ppd* phosphonate biosynthetic genes. Strain names and nucleotide and Ppm accession numbers of gene neighborhoods are: (a) *Lonsdalea britannica* strain 477, NZ_CP023009, WP_094118141; (b) *Zymobacter palmae* strain IAM14233, NZ_AP018933, WP_027705453; (c) *Pectobacterium brasiliense* strain Y60, NZ_JUJP01000005, WP_039555531; (d) *Pectobacterium polaris* strain NIBI01392, NZ_CP017482, WP_095701497; (e) *Pectobacterium polaris* strain F109 KHDHEBDM_11, NZ_RRYS01000011, WP_095993577; (f) *Pectobacterium brasiliense* strain CFIA1001, NZ_JPSM01000013, WP_039282471; (g) *Pectobacterium polaris* strain SS28 SS1, NZ_QESX01000001, WP_109410666; (h) *Pectobacterium peruviense* strain A97-S13-F16, NZ_PYUO01000007, WP_048258525; (i) *Pectobacterium carotovorum* strain S1.16.01.3K, NZ_QZDG01000008, WP_014914040; (j) *Pectobacterium atrosepticum* strain JG10-08, NZ_CP007744, WP_039289843; (k) *Pectobacterium parmentieri* strain IFB5427, NZ_CP027260, WP_012822335; (l) *Pectobacterium wasabiae* CFBP 3304, NZ_CP015750, WP_005971024; (m)*Pectobacterium punjabense* strain SS95, NZ_CP038498, WP_107170185; (n) *Pectobacterium carotovorum* subsp. carotovorum PC1, NC_012917, WP_012773186.

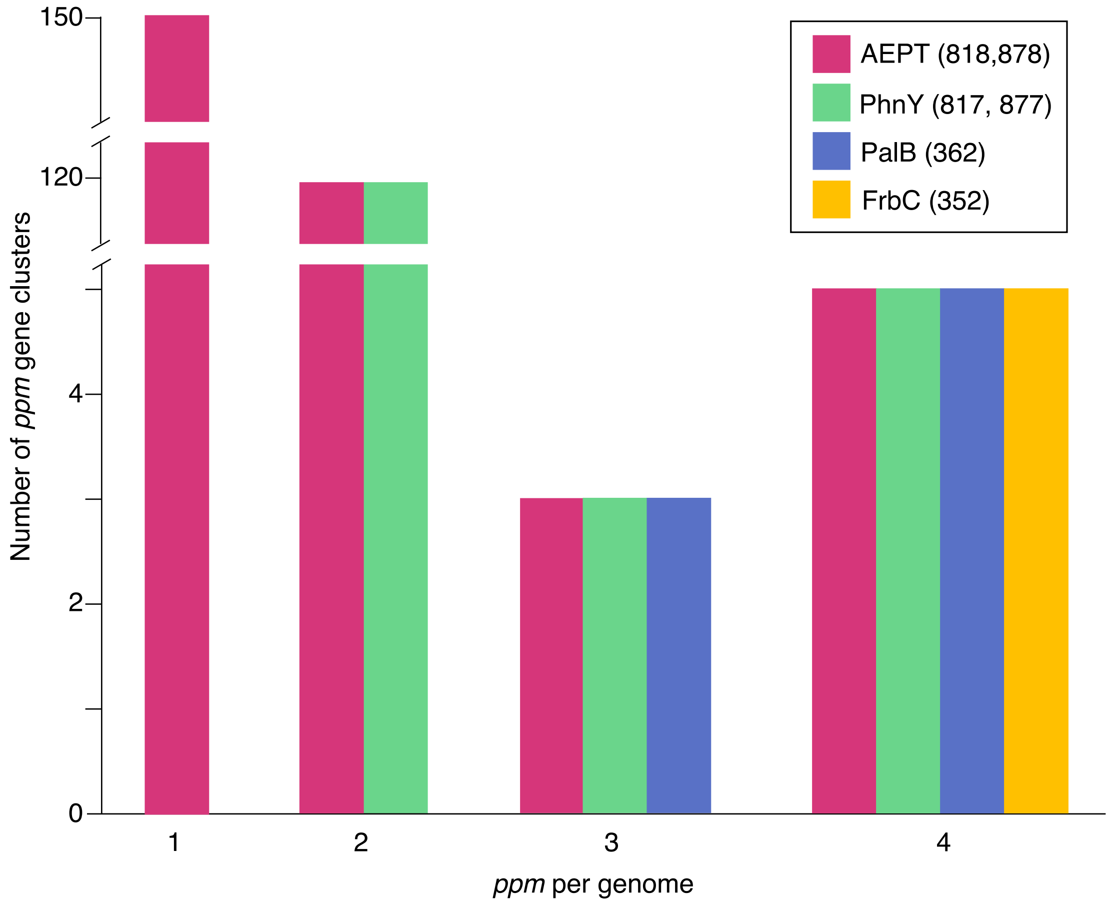


**Figure S7**. Accumulation of phosphonate biosynthetic gene clusters in *Burkholderia*.
